# Alcohol boosts pheromone production in male flies and makes them sexier

**DOI:** 10.1101/2020.08.09.242784

**Authors:** Ian W. Keesey, Georg Doll, Sudeshna Das Chakraborty, Amelie Baschwitz, Marion Lemoine, Martin Kaltenpoth, Aleš Svatoš, Silke Sachse, Markus Knaden, Bill S. Hansson

**Affiliations:** Max Planck Institute for Chemical Ecology, Department of Evolutionary Neuroethology, Hans-Knöll-Straße 8, D-07745 Jena, Germany; Institute of Organismic and Molecular Evolution, Johannes Gutenberg University Mainz, Hanns-Dieter-Hüsch-Weg 15, D-55128 Mainz, Germany; Max Planck Institute for Chemical Ecology, Department of Insect Symbiosis, Hans-Knöll-Straße 8, D-07745 Jena, Germany; Max Planck Institute for Chemical Ecology, Mass Spectrometry/Proteomics Research Group, Hans-Knöll-Straße 8, D-07745 Jena, Germany

## Abstract

The attraction of *Drosophila melanogaster* towards byproducts of alcoholic fermentation, especially ethanol, has been extensively studied ^1–4^. However, the adaptive value of this behavior has not been elucidated. Previous studies have suggested anthropomorphic interpretations of *D. melanogaster* behavior towards alcohols ^5,6^. Here, we instead assert that there exists a simple yet vital biological rationale for alcohol contact and consumption by these insects. We show that exposure to alcohols, especially methanol, results in an immediate amplification of fatty acid ester pheromone levels, which in turn elevates the probability that a male will successfully compete for a female during courtship. We proceed to identify three types of olfactory sensory neurons that detect ethanol and methanol. Moreover, we trace the ensuing neural circuits and reveal their role in controlling both attraction and aversion, where valence is balanced around mating status. Based on our results, we deduce that male flies associate with sources of alcohol as a biological imperative related to reproduction, and we provide an assessment of how and why *D. melanogaster* is associated with alcohol using a sound ecological and natural history approach to this previously enigmatic biological phenomenon.

**One sentence summary:** Flies gain pheromone and courtship advantages with alcohol, but methanol is toxic, thus flies must balance risk versus reward.

---

Mate selection is of paramount importance in all sexually reproducing species, and for many animals, sexual selection drives exaggerated phenotypes. Thus, the coevolution of senders and receivers of sexual signals, such as pheromones, is an ideal ecological context to study adaptation, speciation and animal communication. In *Drosophila melanogaster*, like many insects, the female presumably selects a male based on signals of quality. Moreover, in these instances, it is equally necessary for the males of the species to win any competition against their rivals, as it is prevalent in the *Drosophila* genus for several male suitors to court a single female. Therefore, any competitive advantage in pheromone signaling can have large effects on sexual selection and mating success. In this evolutionary context, we were interested in addressing the role food quality and choice play in competitive advantages that drive preferences in behaviors such as courtship and mate selection.

Species within the genus *Drosophila* are known to be variable in host choice and microbial associations. However, most species of this genus are united by a common behavioral trait, i.e. their preference for vinegar and fermentation odors. Here, given that *D. melanogaster* is a rotten fruit generalist, and that some of the primary byproducts of fruit fermentation are copious amounts of alcohols, we focused on the ecological ramifications of this natural association between alcohol and the fly. In order to examine how flies interact with sources of alcohol, we conducted behavioral studies of attraction (**Figure 1a,b,c**). As shown previously ^4,5,7,8^, male flies were acutely attracted to sources of ethanol. Furthermore, we confirmed that mating status created variability in this attractive behavior ^5^, namely that virgin males were more attracted to sources of ethanol than recently mated males (**Figure 1b**). In addition, we could show that a similar or even stronger increase in attraction was present for methanol (**Figure 1c**), which has not been previously reported. Next we demonstrated that this attraction is attributed to olfaction by testing ORCo anosmic mutants ^9^ (**Figure 1d,e**). Similarly, we addressed alcohol feeding preferences across male physiological states ^5^, and could show that virgin males clearly prefer to consume methanol when compared to mated males (**Figure 1f,g**). Here, all males drank statistically identical volumes, thus the preference for alcohol cannot be explained by increased total consumption (**Figure 1f,g**).

**Figure 1.**
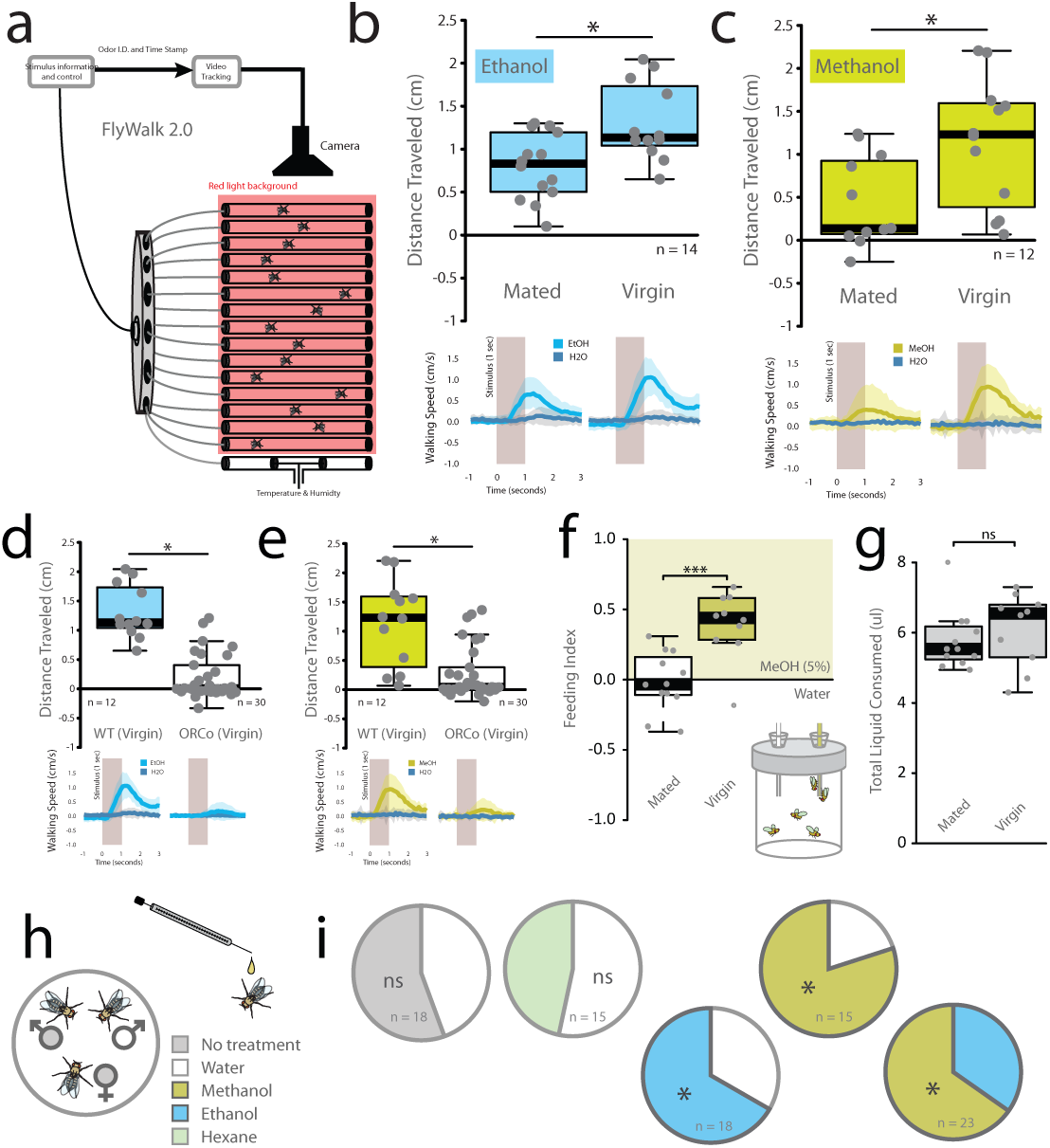
Behavior of Drosophila towards alcohols. (a) Flywalk design schematic for simultaneous video recording and analyses of odor-evoked behavioral responses across 15 flies. (b,c) Flywalk upwind attraction for mated and virgin males relative to water (control), for both ethanol (EtOH; in blue) and methanol (MeOH; in yellow). This matches previous publications for ethanol, though methanol has not been previously examined. Flywalk upwind attraction towards (d) EtOH or (e) MeOH for wildtype and ORCo mutant virgin males. (f) Adults are given a choice between two liquid solutions for consumption, where only virgin males prefer to consume alcohol. (g) Here we show that mated -flies consume as much liquid as the other physiological states, thus it’s a matter of preference for the alcohol by the other physiological states. (h) Competition courtship assays, utilizing two males fighting for access to one virgin female, where shown are percent of total mating for both males. We demonstrate that there is no significant difference when one male is treated with water (control; in white). However, when treated with either methanol or ethanol, that male obtains a significant advantage over the control male. We also show that other organic solvents such as hexane do not have this effect.

Since we had established that virgin and mated males vary by mating status in regards to preference for alcohol, we next wanted to assess any ramifications of alcohol consumption on the associated courtship behaviors (**Figure 1h,i**). We consequently allowed two males to compete for a single female, where we had exposed each male to either a sham or valid treatment. For both methanol and ethanol exposure, those males significantly outperformed their control treated male counterparts (**Figure 1h,i**). Contact with either alcohol thus increased male courtship success dramatically, but with a significantly stronger effect for methanol exposure than ethanol.

We next sought to ascertain which natural host resources produced the highest levels of alcohols during fermentation (**Figure 2a,b**). Here, we could show that citrus fruits, which contain pectin ^10–13^, produced equal or higher amounts of both ethanol and methanol relative to other tested resources (**Figure 2b**). In this context it is important to note that *D. melanogaster* has already been shown to strongly prefer citrus as host fruits ^14^, including studies concerning the ancestral hosts from its African origins ^15^. Next, in order to determine any effects of fermenting host resources on *D. melanogaster* courtship, we repeated our competitive mating assay using male flies exposed to fermented fruit versus standard diet media. Both orange- and banana-exposed males significantly outperformed standard diet males (**Figure 2c,d**). However, treatment with headspace extracts from the same fruits did not result in any mating advantage (**Figure 2e**), where these extracts contain 90-95% of all fruit-derived compounds, but notably do not contain the most volatile odorants, including ethanol and methanol (**Supplementary Figure 1**). In direct comparison, male flies exposed to orange significantly outperformed males from banana (**Figure 2f**), where these fermenting fruits contained roughly equal amounts of ethanol, but a disproportionate amount of methanol (i.e. higher in the orange) (**Figure 2b**). We thus again propose that methanol is more pivotal to this alcohol-related courtship advantage than ethanol.

**Figure 2.**
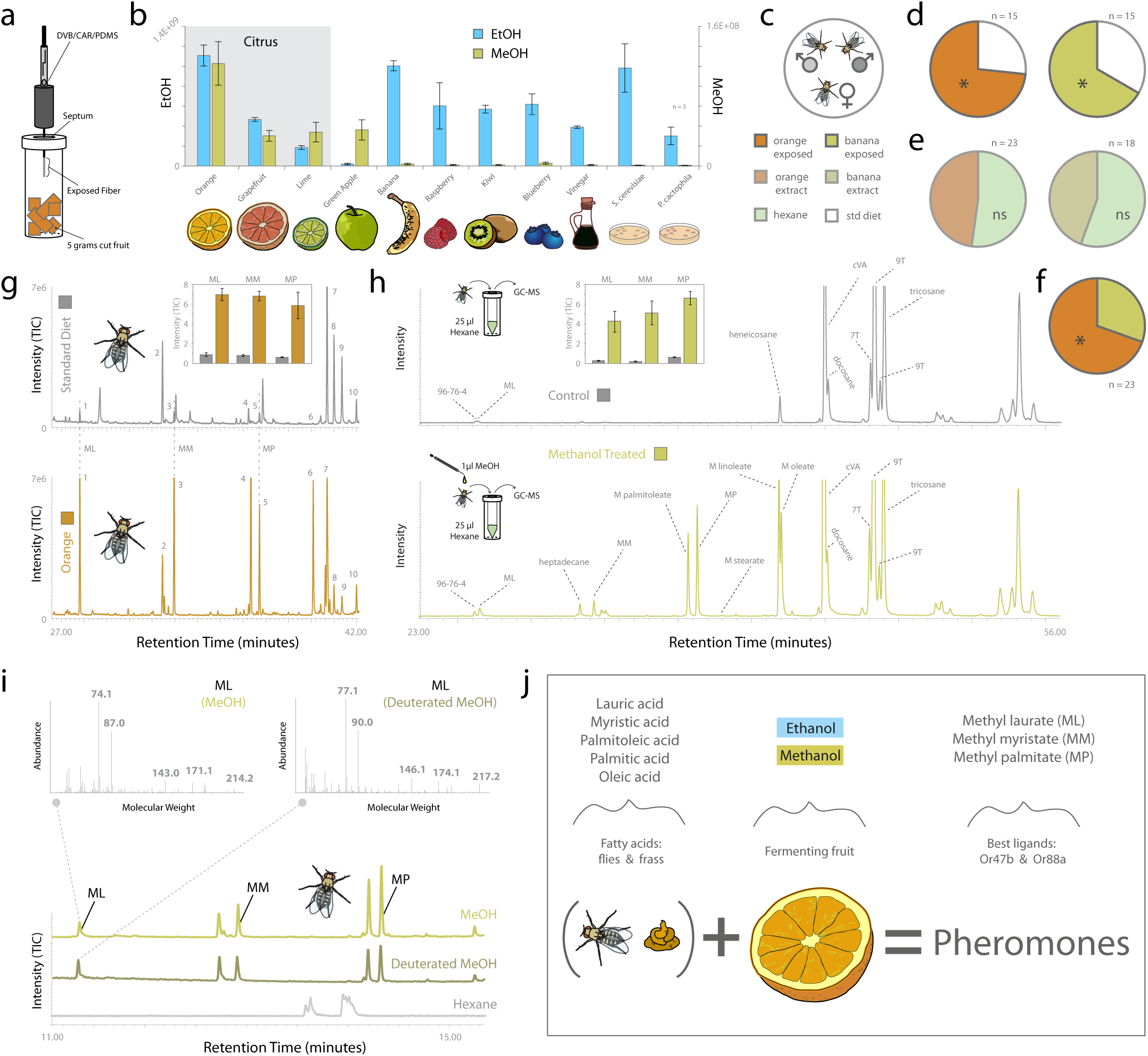
Chemistry and fly pheromone association with alcohols. (a) SPME technique for odorant collection from host substrates, including highly volatile odors such as ethanol and methanol. (b) Quantification of ethanol and methanol content from a variety of natural sources. (c) Competitive courtship paradigm with two males. (d) Orange or banana reared flies perform better than males from water controls. (e) Perfuming the flies with headspace collection of orange and banana does not improve courtship (headspace is specifically missing highly volatile odors like methanol, but has 95% of all other plant chemistry). (f) Orange treated males do better than banana, which correlates strongly with methanol over ethanol content. (g) Thermal desorption (TDU) analyses of flies reared on standard diet with only water (top; in grey); flies reared on fermenting orange (bottom; in orange). Shown are the main pheromone changes following exposure to natural sources of alcohols, including methyl laurate (1), methyl myristate (3), and methyl palmitate (5). (h) Here we assessed the chemical profile of the fly using body wash techniques, both with and without exposure to methanol (bottom, top; respectively). (i) Flies allowed contact with methanol show increase in generation of methyl laurate and other pheromone compounds. To confirm this direct association and hydrogen donation from the alcohol, we use deuterated methanol (d4), where we then show a shift in the mass spectral peaks of all generated pheromones. This confirms that the increase in pheromone (presumably from lauric acid into methyl laurate) is coming directly from the hydrogen or CH3 donation of the alcohol. We also show that this only occurs for the methylated pheromones, and that no other fly body odors, including cVA are deuterated. (j) Ecological overview of alcohol exposure, pheromone increase, and subsequent courtship advantage.

As we had observed increased mating success following exposure to fermenting orange, we then compared the odor profiles of males that had been allowed to contact either citrus or standard diet (**Figure 2g**). In the citrus exposed animals we observed a drastic increase in several established pheromone components, including fatty acid esters known to be involved in both courtship and aggregation behaviors ^16–19^, such as methyl laurate (ML), methyl myristate (MM) and methyl palmitate (MP). In order to examine the involvement of alcohols in the augmented pheromone titers, we pursued experiments with body washes of adult males in several solvents including methanol and ethanol (**Figure 2h; Supplementary Figures 2,3**). Increased production of fatty acid ester pheromones was only observed when the flies had contacted these alcohols, with distinctly larger increases for methanol as opposed to ethanol. This observation matches previous research showing that pheromone-specific olfactory sensory neurons (OSNs) respond more strongly to methyl rather than ethyl esters of these fatty acids ^16,18,19^. To examine any direct role of alcohol in pheromone biosynthesis, we subsequently utilized deuterated isotopologues of methanol (i.e. CD_3_OD instead of CH_3_OH), where these solvents differed only in their isotopic composition, i.e. deuterium replaces hydrogen. This results in deuterated methanol having a distinct shift in mass spectrum, which we could observe during GC-MS analyses. Interestingly, all observed increases in the pheromone profile of the fly following contact with the deuterated methanol resulted in deuterated pheromone compounds (**Figure 2i; Supplementary Figure 3**). Therefore, we assert that contacted alcohols directly donated hydrogen atoms (or -CH_3_ methyl groups) to form fatty acid methyl ester pheromones from fatty acid precursors on the male fly following exposure to methanol (**Figure 2j**). However, putting high purity synthetic fatty acids into alcohol only produced a 1-3% yield of the pheromone compounds (**Supplementary Figure 1c**). Thus, an additional catalyst from the fly must still be required. Subsequent chemical analyses with insects that were devoid of microorganisms (i.e. axenic or conventional) showed that the required catalyst does not originate from a yeast or bacterium, but rather from the fly itself (**Supplementary Figure 4**). However, the exact biosynthetic pathways for pheromone production (i.e. of ML, MM, MP) and the incorporation of methanol hydrogens or methyl groups (-CH_3_) remain elusive.

Given the behavioral motivation of *D. melanogaster* for contact with alcohols, we subsequently examined the mechanism by which the fly detects these odors (**Figure 3**). Whole antenna (EAG) and maxillary palp recordings (EPG) showed that both structures react to ethanol and methanol (**Supplementary Figure 5**). Genetic mutants for the olfactory co-receptor ORCo no longer detected either alcohol, nor produced behavioral attraction, suggesting that odorant receptors (ORs) are necessary for alcohol detection (**Figure 1d,e; Supplementary Figure 5a,b**). We additionally confirmed the OR-mediated detection using mutants deficient in both IR-mediated co-receptors (i.e. Ir8a & Ir25a), which did not show any deficiency in alcohol detection nor attraction (**Supplementary Figure 5c**). Next, we pursued single sensillum recordings (SSR) to ascertain which of the *D. melanogaster* ORs were specifically involved (**Figure 3a-c**). OSNs expressing three different ORs were found to be clearly activated by ethanol and methanol, displaying dose-dependent responses (**Figure 3a-c; Supplementary Figure 5h**). The responses to the alcohols were observed in OSN type ab1A (Or42b) with a stronger affinity to ethanol, and in ab2A (Or59b) and pb1A (Or42a) with stronger responses towards methanol. We also noted a synergy between ethanol exposure and at1A response to cVA (which has been demonstrated previously ^6,20,21^), as well as a possible synergy between at4A responses towards methanol exposure and ML (**Supplementary Figure 5d,e**), although neither alcohol directly activated these trichoid associated OSNs when presented alone. At the OSN level, we did not observe any variation in SSR response based on mating status (mated vs. non-mated males) (**Figure 3d**).

**Figure 3.**
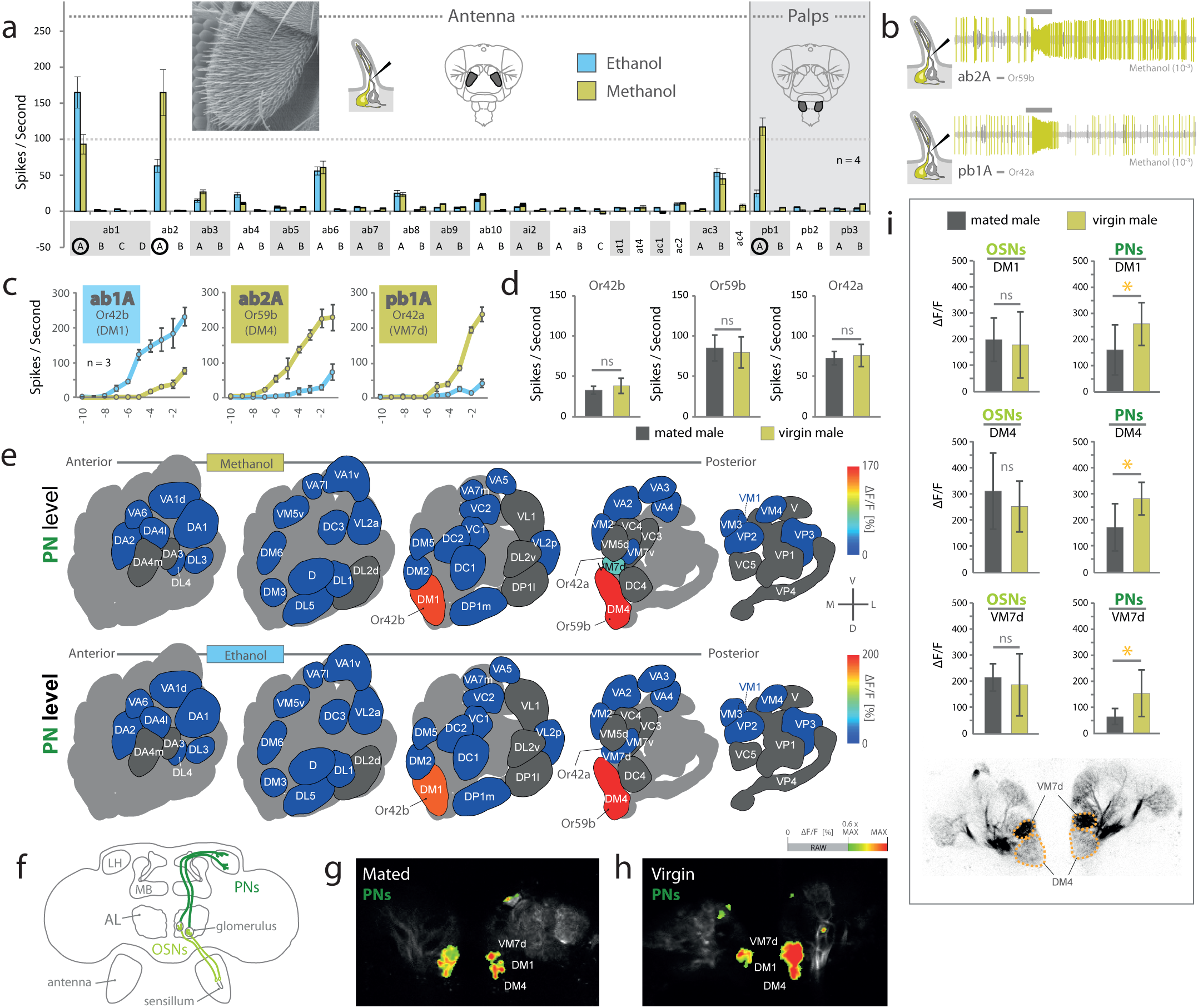
Electrophysiology and pathways for alcohol detection. (a) Entire olfactory screen of all sensillum types across both the antenna and palps of adult wildtype flies using single-sensillum recording (SSR). We identify three sensilla that contain olfactory sensory neurons (OSNs) displaying a response of over 100 spikes per second towards alcohols. (b) Example traces of methanol responses for antennal basiconic 2 (ab2) and palp basiconic 1 (pb1). (c) Highest responding OSN types, with concentration curves for each receptor using both alcohols. ab1A and ab2A show the highest sensitivity for ethanol and methanol, respectively. We also confirm that palp pb1A detects MeOH, though with lower sensitivity than ab2A. (d) SSR responses of mated and virgin males to methanol. (e) Antennal lobe (AL) diagram showing odor-induced calcium responses towards methanol and ethanol (10-3) within the olfactory projection neurons (PNs), again highlighting two primary channels from the antenna for methanol, including DM1 (Or42b; ab1A) and DM4 (Or59b; ab2A), as well as VM7d (Or42a; pb1A) from the palps, which is then supported by our SSR data from the periphery. (f) Diagram of D. melanogaster brain, highlighting OSN and PN circuits. (g-h) Examples of odor-induced false color-coded raw images for PNs from mated and virgin males responding to methanol. (i) Odor-induced fluorescent activity towards methanol, as recorded from the OSNs input and PNs output that exit the AL and extend towards the mushroom body (MB) and lateral horn (LH). Here we observe significant differences in calcium responses associated with physiological state (i.e. mating status) within these PN types.

Moving from the periphery into the brain, using optical imaging of calcium dynamics in OSNs and PNs, we found that mainly two glomeruli were activated by methanol and ethanol after stimulation of OSNs on the antenna (**Figure 3e; Supplementary Figure 5f,g**), namely DM1 (innervated by OSN ab1A expressing Or42b) and DM4 (ab2A; Or59b). In correspondence with the SSR responses of the two innervating OSN types, the two glomeruli were preferentially tuned towards ethanol (DM1) or methanol (DM4), respectively (**Figure 3a,c; Supplementary Figure 5f,g**). After stimulation of OSNs on the palps, a third glomerulus, VM7d (pb1A; Or42a), was activated. The affinity was clearly higher towards methanol than ethanol, though with lower sensitivity than that observed in DM4 (**Figure 3a,c,e**). Interestingly, all three glomeruli that detect the alcohols are close neighbors within the AL, suggesting perhaps a similar evolutionary origin ^22^ (**Figure 3e**).

In order to revisit the role of mating status, which produced variation in behavioral attraction towards sources of alcohol (**Figure 1**), we next examined at what neural level we could observe such variance within the brain. Here, mated and virgin males showed no difference in sensitivity at the level of OSNs, neither in SSR nor in optical imaging of OSN responses in the AL (**Figure 3d,e,f,i; Supplementary Figure 5f,g,i**). However, virgin males did display a significantly higher sensitivity towards methanol when we imaged calcium dynamics in projection neurons (PNs), which represent the output elements of the AL (**Figure 3f-i; Supplementary Figure 5k**). A potential neural correlate to the state-dependent increase in attraction towards alcohol thus appeared in neural connections towards higher brain structures (i.e. mushroom body (MB) and lateral horn (LH)) and after processing within the AL. This mating status-dependent increase was specific to alcohol circuits and did not appear within food-related neural pathways (**Supplementary Figure 5j; DM2**).

Many of the previously characterized dedicated circuits for olfactory behavior utilize only a single neural pathway ^7,14,23^, thus we next wanted to explore why these flies possess three separate pathways for alcohol detection. We revisited attraction paradigms using transgenic flies, where we tested flies with individually silenced OSNs expressing one of the three main alcohol-detecting receptors (**Figure 4a-c**). While each silenced OR on the antenna (i.e. Or42b, Or59b) exhibited a pronounced loss of attraction compared to parental controls, the third silenced OR (Or42a), located on the palps, resulted in a significant increase in attraction. This suggested that two circuits relate to attraction to alcohols, while the third mediates aversion towards methanol. Using single PN labeling and reconstruction in combination with already published material ^24–26^, we next assessed the innervation patterns from the AL towards the MB and LH for all three alcohol-related circuits (**Figure 4d-f; Supplementary Figure 6**). Two alcohol pathways displayed a high degree of overlap (i.e. DM1 and DM4, both innervating from the antenna, and coding for alcohol attraction) with major branches in the MB and overlapping innervation areas in the LH. However, the third (emerging from VM7d; innervating from the palp and coding for methanol aversion) displayed only minor branches within the MB and a clearly separate innervation area within the LH (**Figure 4e**). Moreover, the innervation pattern of this pathway (VM7d) matched well with other known aversive circuits in the MB and LH (**Figure 4f; Supplementary Figure 6b**). This difference in innervation pattern was not due to VM7d originating from the palps, as opposed to the antenna (**Supplementary Figure 6e**). Reconstructions also revealed that while the two attractive alcohol circuits possessed only a single PN each, the VM7d glomerulus sent three PNs to higher brain areas ^23^. Such a high PN to OSN ratio has been proposed as another characteristic of labeled line circuits, and is also common to several aversive neural pathways ^23^.

**Figure 4.**
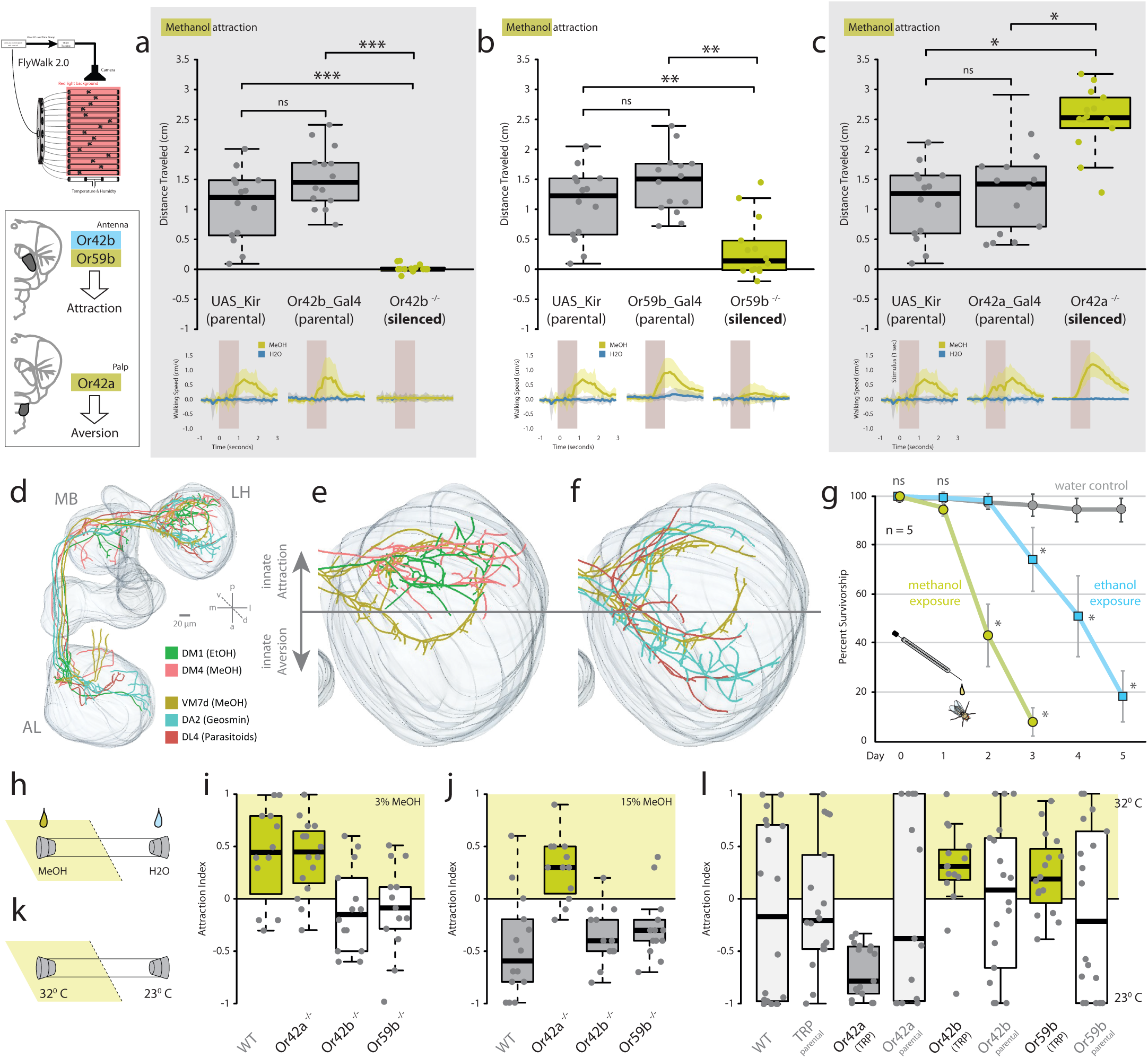
Neural circuitry for behavioral attraction and aversion towards alcohol. Flywalk behavior and upwind attraction for wildtype and single-neuron silenced virgin males relative to water (control) for methanol (MeOH; in yellow). (a) Methanol attraction for parental lines and loss-of-function silenced neuron for Or42b (ab1A; DM1) (b) Or59b (ab2A; DM4) (c) Or42a (pb1A; VM7d) (d) Complete neural circuit reconstruction of projection neurons (PNs) for DM1, DM4 and VM7d (i.e. all alcohol detecting circuits), in comparison to two known aversive circuits emerging from DA2 (geosmin-detection) and DL4 (parasitoid-detection). (e) Neural circuit reconstruction of PNs extending into the mushroom body (MB) and the lateral horn (LH) for DM1, DM4 and VM7d (i.e. all alcohol detecting circuits). We highlight that our two circuits related to attraction (DM1, DM4) match previous studies suggesting upper portions of the LH relate to innate attraction, while the PNs from our potentially aversive circuit map to lower sections of the LH (VM7d). (f) Overlay of our potentially aversive circuit (VM7d) with neural circuits from PNs for two well-studied labeled lines for aversion (DA2 and DL4). (g) Survivorship during repeated exposure to alcohols. (h,k) Behavioral schematics for high and low concentration behaviors within tube assays, as well as for temperature activation of the three different single-receptor silenced lines. (i) Behavioral responses of flies towards low concentration of methanol (3% MeOH). (j) Behavioral response of flies towards high concentration of methanol (15% MeOH). (l) Responses of thermally activated single circuits and genetic controls.

Why would *D. melanogaster* maintain pathways for both attraction and aversion towards the same odorants? In this case, we identify that both alcohols are toxic at higher concentrations, with methanol more so than ethanol, and thus contact bears an inherent risk for flies (**Figure 4g**). We hypothesized that alcohols might therefore be attractive at low concentrations, but aversive at higher ones. Again using single receptor loss of function mutants, we tested attraction at both low and high, ecologically relevant concentrations of methanol (**Figure 4h,i,j**). In these experiments, we observed that two receptors are necessary for attraction to low alcohol levels (Or42b, Or59b), while aversion to high alcohol content is dictated by a single receptor (Or42a). Furthermore, we artificially activated these three circuits independently and without odor stimulation using thermogenetics, and could again surmise that while two receptors drive attraction, the third (Or42a) was sufficient to drive aversion when this odorant receptor was activated by heat alone (**Figure 4k,l**). This is similar to ecologically relevant concentrations of CO_2_, where two neural pathways are weighed against each other in order to control behavioral valence ^7^. Crosstalk has also been shown to occur between information regarding different types of odorants ^20,27,28^, and may play a larger role in physiological state-dependent coding, as well as in determining feeding and oviposition preferences across the *Drosophila* genus, for example, in divergent preferences for different stages of host fermentation ^29–31^.

In summary, we show that contact with alcohols produces an escalation in the release of aggregation and courtship pheromones and subsequently leads to increased male mating success. We further assert that this pheromone increase is potentially a signal of male fitness and male quality. As while methanol itself is a highly volatile and fleeting odor, contact with this alcohol contributes directly to an increased production of a more environmentally stable pheromone signal (i.e. methyl laurate). This thereby advertises the ability of a male to successfully find and utilize optimal stages of host decay, or signals the ability of the male to withstand high concentrations of otherwise toxic alcohols. We thus propose that this makes males more attractive for females that are seeking good genes for their progeny. Moreover, the increased attraction to alcohol by virgin males appears to give a direct advantage in subsequent courtship, as opposed to depression or substance abuse interpretations of this behavior ^5,6^. However, contact with alcohols, especially methanol, is inherently dangerous, as it is toxic. Here we show that the separate positive and negative methanol-coding pathways provide the neural substrate necessary for such a behavioral decision and risk assessment by the fly. Thus, adult *D. melanogaster* must carefully examine benefits versus costs for alcohol exposure, where the rewards might sometimes outweigh the risks, especially for virgin males seeking to compete successfully for a mate.

## Acknowledgements

This research was made possible by the Max Planck Society (Max Planck Gesellschaft). ML and MKa acknowledge funding from the German Research Foundation (DFG KA2846/5-1) as part of the consortium project FOR 2682 ‘Seasonal temperature acclimation in Drosophila’. Wild-type flies were obtained from the San Diego Drosophila Species Stock Center (now The National Drosophila Species Stock Center, Cornell University), while mutant and silenced lines were obtained from the Bloomington Stock Center. We express our gratitude to K. Weniger, S. Trautheim, M. Niebergall and D. Veit for their technical support, expertise and guidance at MPI-CE, and T. Engl for assistance with chemical analytics at the University of Mainz. We also thank Richard Benton as well as Angela Douglas and her team at Cornell University for their support and guidance in regards to this publication.

## Author Contributions

IWK generated the original hypotheses and concept for this study, with support from SDC, SS, MKn and BSH. Bioassays and electrophysiology were conducted by IWK. Chemical analyses were performed by IWK in association with AS, as were the measurements and assessments of pheromones and deuterated components. IWK and GD collected and analyzed courtship videos and other behavioral data. Analyses of olfactory sensory neurons, antennal lobe, and projection neurons were all conducted by SDC, AB and IWK, under the supervision of SS. ML, MKa, and IWK performed the experiments pertaining to axenic or conventional flies. All diagrams, illustrations and figures were created by IWK, as was the original written manuscript. Each author subsequently improved the final version of the writing and figures, as well as assisted with any revisions towards the final publication.

## Data Availability

All data supporting the findings of this study, including methodology examples, raw images, raw data and z-stack scans, molecular sequences, statistical assessments, as well as transgenic insect information are all available with the online version of this publication via Edmond, the Open Access Data Repository of the Max Planck Society.

## Methods

### Fly stocks and insect rearing

Transgenic fly lines were obtained from the Bloomington *Drosophila* stock center (http://flystocks.bio.indiana.edu/), and diets for rearing are included within the supplementary information. Unless otherwise noted, all fly stocks were maintained on standard diet (normal food) or yeast food (i.e. axenic assays) at 25 °C with a 12 h light/dark cycle in 70% humidity. Stock population density and size was controlled by using 20–25 females per vial. Stocks were maintained according to previous publications ^33^, and for all behavioral experiments we used 2-7 day old male flies. For mating status experiments, we collected males as virgins and then later divided them into three cohorts. We gave one group of males access to intact, virgin females, and allowed them to mate and complete copulation within one hour prior to the onset of behavioral experiments (e.g. mated). A second group was not ever exposed to any females (i.e. virgin). Thus, we controlled for age of the males, with the only differences occurring within the one hour window of time prior to onset of the behavioral experiments. We did not consistently observe a significant difference in behaviors for sexually frustrated males, i.e. virgin males allowed access to a headless female, as compared to non-frustrated virgins, and in general, alcohol attraction was not significantly different between frustrated and non-frustrated virgin males. As such, we have focused solely on the behavior and neurological changes between mated and virgin males.

### Flywalk attraction assay

Behavioral experiments were performed in the Flywalk paradigm as previously described ^34,35^, except with male flies starved for only 1-3 hours before the start of the experiments. In short, 15 individual flies were placed in 15 glass tubes (inner diameter 0.8 cm). Glass tubes were aligned in parallel, and flies were continuously monitored by an overhead camera (SONY EVI, Sony Corporation, Japan) under red-light conditions (λ > 630 nm) (Figure 1A). During the experiment, flies were continuously exposed to a humidified airflow of 20 cm per second (70% relative humidity; 20°C). We presented flies repeatedly with pulses of different olfactory stimuli at an interstimulus interval of 90 seconds. Stimuli were added to the continuous airstream and thus traveled through the glass tubes at a constant speed. Odor stimulation was performed with a multicomponent stimulus device described elsewhere ^34,35^. In summary, 100 μl of an odor was prepared in 200 μl PCR tubes without cap, where these tubes were placed in odor containers made of polyetheretherketone thermoplastic. These odor containers were tightly sealed and connected to the stimulus device via ball-stop check valves.

These valves only allow uni-directional airflow through the odor-saturated headspace. Odor stimulation was achieved by switching an airflow otherwise passing through an empty vial (compensatory airflow) to the odor-containing vial. Odor pulses were 500 ms in duration, again at an interstimulus interval of 90 seconds. Different stimuli were presented in pseudorandomized order (random block design) to avoid odor-sequence artifacts. A control stimulation was always included in the randomized block, which consisted of only the humidified water. Please see supplementary videos and the included diagrams for more complete descriptions of these behavioral assays.

### Café feeding assay

All tested flies were 2 to 7 days old, which included only males, and flies were starved beforehand for 18–20 hours with constant access to water. Flies then were cooled for 5 min at −20 °C to assist in their transfer (without CO_2_) to plastic vial arenas (Figure 1D). Basic feeding solutions consisted of water with 5% sucrose, as well as 5% MeOH (treatment) or without (control). The capillary feeder (CAFÉ) assays utilized glass micropipettes with liquid media that were filled by capillary action, and then inserted through pipette tips into the container holding the adult flies ^17^. Each assay had both control and treatment options for feeding. The volume consumed from each capillary was measured after a set duration. The feeding index was calculated as (treatment – control) / total volume consumed.

### Courtship assay

For the analysis of courtship behavior, the adult flies were collected as pupae and moved into single vials (using a wet paint brush), then later identified by sex after subsequent eclosion. Adults were kept in these single vials for 2 – 7 days after eclosion with access to food and water. Temperature controlled chambers were used for courtship conditions. In these behavioral assays, we first aspirated a female fly into the tiny chamber, and secured the chamber with a clear cover slide. Next, two male flies were introduced into the same chamber, and video recording was initiated. The flies were recorded under white light illumination for 10 – 30 minutes. If no initiation of courtship was observed after 30 minutes, then videos were halted and new flies were introduced as a novel pair. Videos of successful courtship and copulation were analyzed with BORIS (http://www.boris.unito.it/). More information about the arena was described previously ^36^. In assays with two males competing for a single female, one male was always marked with fluorescent powder. In order to avoid any potential bias from the powder, males were allowed to clean themselves for 12 hours before experiments, and powder was given to either control or treated flies randomly. This fluorescent powder was easily observed under UV light, and thus we could confirm the identity of each male after copulation success. Mating and courtship behaviors were recorded by video and then analyzed using BORIS software (http://www.boris.unito.it/). We visualized the data as the percent copulation success for each of the two males (i.e. control vs. treatment). Here we performed 15 to 23 replicates of each male competition courtship assay (Figure 1G; Figure 2C-F). Treatment males received a single droplet of 1 µl treatment while under CO_2_ anesthesia, and control males received the same but only water, then all flies were allowed 1-2 hours of recovery before the onset of behavioral trials.

### Odor collections, SSR, GC-MS, and TDU-GC-MS

All synthetic odorants that were tested in this publication were acquired from commercial sources (Sigma, www.sigmaaldrich.com; Bedoukian, www.bedoukian.com) and were of the highest purity available. Stimuli preparation and delivery for behavioral experiments followed previously established procedures, and collection of volatile and non-volatile compounds was carried out according to standard procedures ^17,18,31^. GC-MS (HP5 and HP-Innowax; liquid samples) and TDU-GC-MS analyses (HP5 and HP-Innowax; single fly, solid samples) were performed on all odor collections and insect body washes as described previously ^16^. The NIST mass-spectral library identifications were confirmed with chemical standards where available, and the internal standard bromodecane was utilized for quantification and statistical comparisons between analyzed pheromone samples. Single sensillum recordings (SSR) experiments were conducted as described previously ^16,18,29^. Adult flies were immobilized in pipette tips, and the third antennal segment or the palps were placed in a stable position onto a glass coverslip. Sensilla were localized under a microscope (BX51WI; Olympus) at 100x magnification, and the extracellular signals originating from the OSNs were measured by inserting a tungsten wire electrode into the base of a single sensillum. The reference electrode was inserted into the compound eye. Signals were amplified (10x; Syntech Universal AC/DC Probe; Syntech), sampled (10,667.0 samples/second), and filtered (30 – 3,000 Hz, with 50/60 Hz suppression) via USB-IDAC4 computer connection (Syntech). Action potentials were extracted using Auto Spike 32 software (Syntech; v3.7). Neuron activities were recorded for 10 seconds, with a stimulation duration of 0.5 seconds. Responses from individual olfactory sensory neurons were calculated as the increase (or decrease) in the action potential frequency (spikes per second) relative to the pre-stimulus spike frequency.

Concentrations of alcohols were calculated from fermented samples. Five grams of each organic fruit type was collected into sealed glass vials, and allowed to ferment with minimal airflow for 2 days. Solid phase microextraction (SPME) using triple action fibers (DVB/Carb/PDMS) provided collection of all volatile odorants from each sample, replicated three times. Fibers were cleaned between collections using manufacturer recommended heating protocols under clean helium streams. Peak area for each alcohol was calculated, and averaged across all three replicates. Alcohol content from both yeasts were measured from 5 mL liquid cultures, using identical glass vials, SPME fiber and volumes as solid host materials. Methanol and ethanol are shown at different scales (Figure 2B), with ethanol 10x times higher.

### Deuterated solvents

Methanol and ethanol of the highest possible purity was used wherever possible (>99.5% purity, reinst, CP43.1 (MeOH), 5054.2 (EtOH); Carl Roth GmbH). For deuterated solvents, we used previously published guidelines ^37^ to identify shifted mass spectral peaks between methanol and deuterated methanol treated adult male flies (D4, 100Atom%D, CAS: 811-98-3; Carl Roth GmbH).

### Calcium imaging (OSN, AL, PN)

We dissected flies for optical imaging according to standard protocol ^21,38^. Flies were briefly immobilized on ice and then mounted onto a custom-made stage. Protemp II composite (3M ESPE) was used to fix each head. We bent the anterior part of the fly’s head with fine gold wire, and a small plastic plate having a round window was placed on top. We sealed the head with that plate using two-component silicone (Kwik Sil), leaving the center part open to make a cut. The cuticle between the eyes and the ocelli was cut under saline solution (130 mM NaCl, 5 mM KCl, 2 mM MgCl_2_, 2 mM CaCl_2_, 36 mM saccharose, 5 mM Hepes, 1 M NaOH, pH 7.3). Then the cuticle was either bent forward and fixed to the silicon, or removed. After cleaning the fatty tissues and trachea, we were able to visualize the antennal lobes (AL).

We used a Till Photonic imaging system with an upright Olympus microscope (BX51WI) and a 20x Olympus objective (XLUM Plan FL 20x / 0.95 W), as described previously for the functional imaging ^20^. Among the odorants, methanol (99.5% from Sigma) was diluted in double distilled water (ddH_2_0) to make concentrations of 10^−3^, 10^−4^, and 10^−5^, 10^−6^, and ethanol was diluted in ddH_2_0 to make the same concentrations. Six microliters of these dilutions were pipetted on a filter paper (∼1 cm2; Whatman), which was placed in Pasteur pipettes. We used filter papers with ddH_2_0 water control alone, as blanks. A stimulus controller (Stimulus Controller CS-55; Syntech) was used for odor application. Continuous airflow (1 Liter/min) and pulses of odor (0.1 Liter/min) were directed through an acrylic glass tube to the antenna of the fly. Odor stimuli were injected into this airstream after 2 seconds for a duration of 2 seconds. The recording frequency during imaging was 4 Hz with 40 frames (i.e., 10 seconds) in total. Each odor was measured only once in each animal and the odor stimulation sequence was delivered from low concentration to high concentration for each experiment. However, not all concentrations could always be measured in all animals, as some insects died during testing. Therefore, the number of animals for each concentration might differ, but sample size is given in each plot, and raw data are available with the online version of this publication. The interstimulus interval was at least 60 seconds to avoid any effects of adaptation or habituation. To test whether the odor responses were reproducible from trial to trial, we measured repeated stimuli in single animals and observed that all three consecutive repetitions induced a similar response.

We analyzed data with custom-written IDL 6.4 software (ITT Visual Information Solutions). Manual movement correction and bleach corrections were followed by the calculation of relative fluorescence changes (ΔF/F) from the background. The glomeruli were identified according to previous publications ^23^. The ΔF/F of all 40 frames was imported to an Excel file. The responses from frames 10–18 were averaged for the glomerulus of interest for all treatments. Wilcoxon matched paired test was used for all statistical analyses of the imaging data.

### Neural 3D reconstruction, tracing and mapping

For in vivo photoactivation experiments, 1–6 day old flies (genotype: END1-2,UAS-C3PA;MZ699-GAL4) were dissected, and the tracts of the salivary glands were cut to prevent movement. Photoactivation was accomplished via continuous illumination with 760 nm for 15–25 minutes. After a 5-minute break to permit full diffusion of the photoconverted molecules, 925 nm Z-stacks of the whole brain were acquired and subsequently used for neuronal 3D-reconstruction. For all 3D reconstructions, we used the segmentation software AMIRA 4.1.1 & 5.3.3 (FEI Visualization Sciences Group, Burlington, MA). Neurons of different individuals were embedded into the reference brain using a label-field registration as previously described ^39^. Briefly, segmented labels of brain neuropils (antennal lobe: AL, mushroom body: MB, lateral horn: LH) were registered onto a reference brain image using affine registration followed by elastic warping. In a second step, the calculated transformation matrix was applied to the respective neuron morphology that was then aligned to the reference brain image ^40^.

Photoactivation and transection procedures as well as image acquisition following immunohistochemistry were accomplished with a 2-photon confocal laser scanning microscope (2PCLSM; Zeiss LSM 710 NLO) equipped with a 40x (W Plan-Apochromat 40x/1.0 DIC M27; Zeiss) or 20x (W N-Achroplan 20x/0.5 M27; Zeiss). The 2PCLSM was placed on a smart table UT2 (Newport Corporation, Irvine, CA, USA) and equipped with an infrared Chameleon Ultra diode-pumped laser (Coherent, Santa Clara, CA, USA). Z-stacks were performed with argon 488 nm and heliumneon 543 nm laser or the Chameleon Laser 925 nm (BP500-550 for G-CaMP and LP555 for DsRed/tdTomato) and had a resolution of 1024 or 512 square pixels. The maximum step size for immuno-preparations or single neuron projections was 1 µm, and for AL reconstructions 2 µm.

Reconstructions of single neurons were also compared to previously published, online datasets, including those obtained from Virtual Fly Brain (VFB; https://v2.virtualflybrain.org/). Virtual Fly Brain uses an ontological model of *Drosophila melanogaster* anatomy written in OWL2 and based on the *Drosophila* literature. This contains detailed information about gross neuroanatomy, neuron classes, as well as the relationships between them. Underlying each neuroanatomy query is a query of this ontology in OWL-DL. Queries of phenotype and expression first utilize the large volume of expression and phenotype data available from FlyBase (https://flybase.org/), and then annotate using the *Drosophila* anatomy ontology. Each expression or phenotype query starts with a query of the anatomy ontology for terms appropriate to the chosen region. The output of this query is then used as input for a query of the FlyBase database for expression or phenotype annotated using these terms. Images in the online viewer are delivered as a series of tiles covering only the visible area in the browser window. The tiles are produced from a compound 3D Woolz object (https://github.com/ma-tech/Woolz), representing the overall structure and individual color painted domains. Here we use the neural tracings of following projection neurons: DM1 (VFB_00101219), DM4 (VFB_00101234), Vm7d (VFB_00101137), DA2 (VFB_00101260), DL4 (VFB_00101235), and VC2 (VFB_00101165).

### Tube assays (with odorants or temperature gradients)

Male flies were collected shortly after emergence, as described previously for virgins. We kept all flies at 22-23^0^ Celsius in 70% humidity for 24 hours within the behavioral chambers, prior to onset of their use in experiments. Single adults were placed into tubes described from Flywalk assays, but without dynamic headspace. Each end of the tube was sealed with medical-grade cotton, and an aliquot of 50 µl of water (control), or 50 µl of alcohol diluted in water (treatment) was added to opposite sides of the tube. We allowed flies to acclimate for 10 minutes without disturbance, and then we recorded their resting position within the tube after an additional five minutes of observation. We utilized the fly position (in cm) from the control side to generate an attraction index, with higher numbers indicating proximity to the treatment. For heat sensitive loss-of-function or heat activation lines, we used a similar assay consisting of the same glass tubes. However, instead of odors, one side of the glass tube was heated to 30-35^0^ Celsius (treatment) while the other side remained at ambient temperature (20-24^0^ Celsius). A Bosch PTD 1 laser thermometer (Robert Bosch Power Tools GmbH; Stuttgart, Germany) recorded the temperatures of both sides of the glass tubes at the onset of each experiment, where three glass tubes and fly replicates were usually run in parallel.

### Toxicity and alcohol exposure

We collected male flies as virgins after eclosion, and sorted them into cohorts of 20 adults per rearing vial. Each cohort received either a high dose of ethanol or methanol twice per day (morning and evening) for one week. Each day, cohorts were anesthetized with CO_2_ and then each single fly was given a 1 µl droplet of alcohol, and then placed back into the standard rearing vials with access to regular food substrates. We recorded survivorship each day, and present the data as an average of five replicates of 20 flies.

### Generation of axenic and conventional flies

Axenic flies were generated using modifications towards previously established protocols ^41^, in this case to collect first instar larvae, and these methods are further described herein. Adult females were mated and then allowed to oviposit their eggs onto apple-juice plates. Before egg transfer, we cleaned tools and the new fly food for rearing with ethanol and with exposure to UV light for 15 minutes inside a sterile bench for starvation media (i.e. apple-juice plates), where the eggs were then transferred after the bleaching step. The apple-juice plates where the flies laid eggs were not sterilized before use, but the media was autoclaved and poured in the clean bench. After oviposition, we rinsed the apple-juice plates with PBS 1x, gently brushed the surface, and then poured the contents into two sieves for egg collection. The collection contents from both sieves were washed three times with PBST (0.1 % triton X), re-suspended for 35 seconds, and then rinsed again with distilled water. Next, in the clean bench, the eggs were washed in 3% bleach solution by transferring over another sieve into a sterilized container, and subsequently cleaned again with distilled water. All eggs, including axenic and conventional, were stored separately for one day in an incubator at 22^0^C with 60% humidity until larvae could emerge and be collected. Here again, the axenic fly larvae were kept to food media that was previously autoclaved and then poured into tubes in the clean bench. We exposed these food vials to UV light and we kept them in sterile closed containers until use.

**Supplementary Figure 1.**
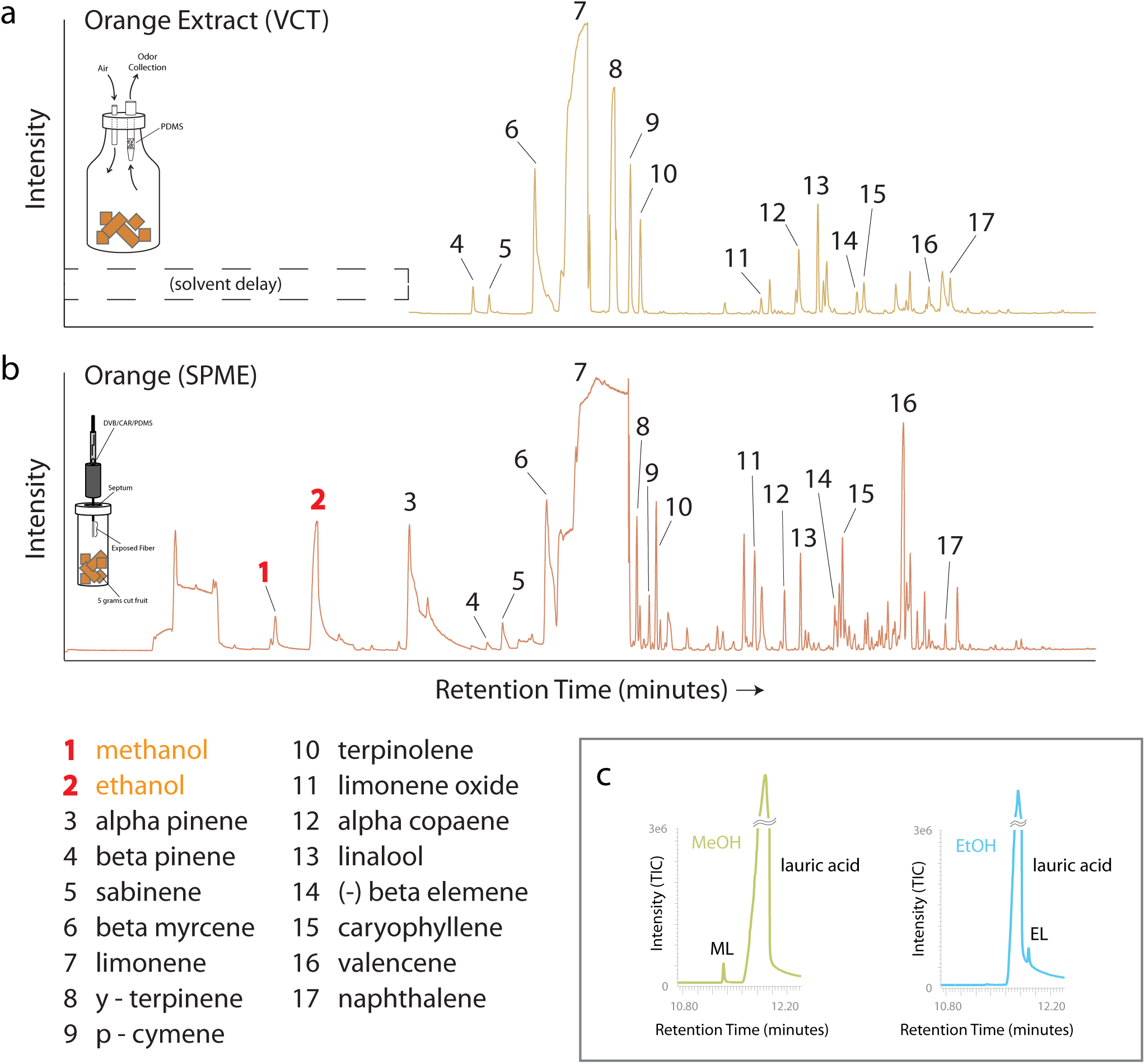
Odor samples from fermenting oranges. In order to examine the roles of the numerous compounds generated from orange fruit material potentially involved in the increased production of pheromone components after fly exposure to this fermenting fruit, we collected samples in two ways. Shown are the GC-MS total ion chromatograms (TICs) from each collection method using identical host plant materials. (a) The headspace from sliced oranges was collected using volatile collection traps (VCTs; PDMS absorbent), and then eluted using hexane solvent. We subsequently used these liquid samples to perfume male flies for courtship experiments (Figure 2E), and in addition, these headspace collections were run across a GC-MS for further analyses. (b) The headspace of sliced oranges was collected using solid phase micro extraction (SPME), and then analyzed by injection into the same GC-MS. Here we note nearly identical odor identities collected with these two sampling techniques. However, importantly, VCT collections did not contain highly volatile odors when compared to SPME (which are lost during sample collection), and this would include the loss of both alcohols. Thus greater than 90% of odors are still present in courtship perfume trials (Figure 2E) but fail to create a courtship advantage without the presence of alcohols. (c) Putting lauric acid in alcohols only gives 1-3% yield of pheromone (therefore missing a potential catalyst from fly or microorganism).

**Supplementary Figure 2.**
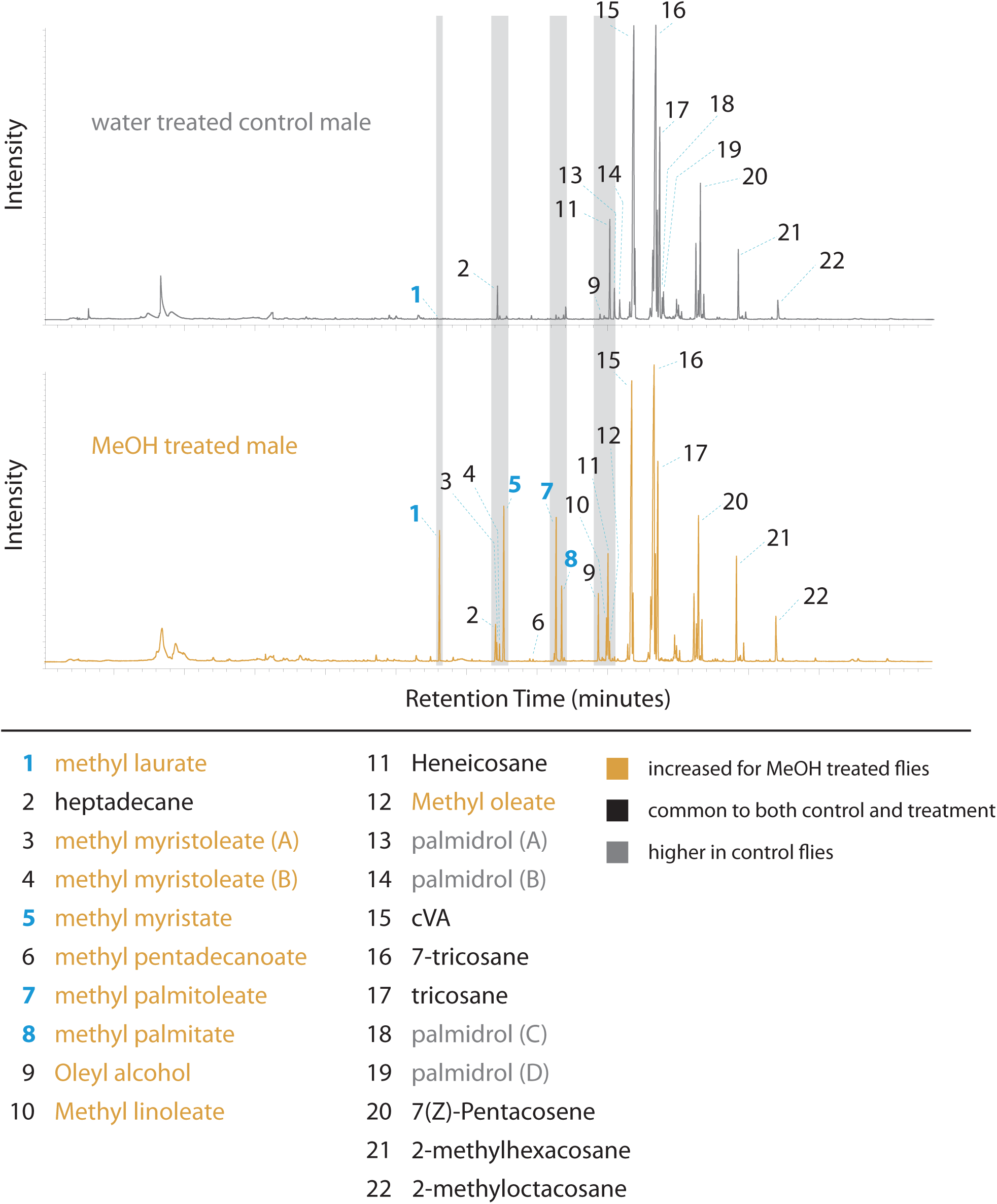
TDU-GC-MS chemical analyses of adult males. In order to assess the chemical profile of adult males in the absence of solvent injection, we utilized a single, live fly thermal desorption unit (TDU). Here we either pre-treated adults with a 1ul drop of water (top; grey) or a 1ul droplet of 10% methanol (MeOH; diluted in water; bottom, orange). We then thermally desorbed live flies into the GC-MS. Here we noted increases for several, but not all body odors emanating from the insect. For example, we did not observe any changes to cuticular hydrocarbons (CHCs) or for cVA (which is produced by the male accessory glands). However, we again note strong increases for several fatty acid methyl esters, including behaviorally active ML, MM, and MP (peaks 1, 5, and 8).

**Supplementary Figure 3.**
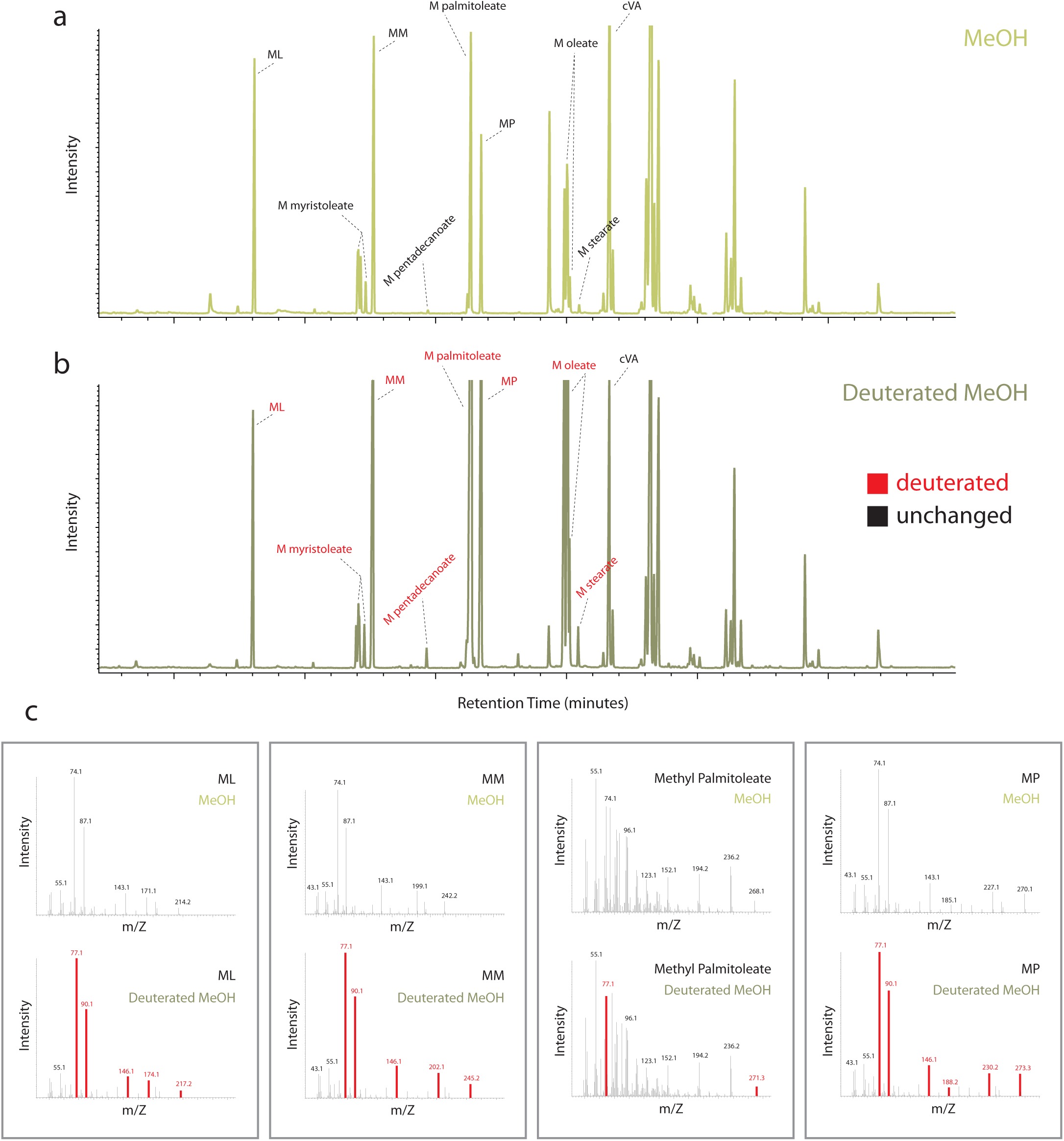
Use of deuterated methanol (d4) to assess pheromone increases. Shown are examples of two GC-MS total ion chromatograms (TICs) of a single male drosophila exposed to alcohol, with methanol at top (a), and deuterated methanol at bottom (b). Here we observe nearly identical TICs between these two treatments; however, several pheromone compounds have their molecular weight shifted in the case of the deuterated methanol treatment (bottom). This would include methyl laurate (ML), methyl myristate (MM), methyl palmitoleate (M palmitoleate), as well as methyl palmitate (MP). We did not observe any changes to other known pheromones, such as cVA, or cuticular hydrocarbons, such as 7-tricosane or 9-tricosane. Thus, this change in molecular weight following deuterated methanol exposure is limited to only those compounds that were increased following alcohol exposure. (c) Shown are the mass spectra of pheromones from methanol (top) and deuterated methanol exposed male Drosophila (bottom). In each case, we document a consistent shift in molecular weight for these fatty acid pheromones when the fly contacts deuterated alcohols, suggesting the pheromones are produced directly by the alcohol’s donation of three deuterated hydrogen atoms. For example, from mass 74.1 to 77.1, or 87.1 to 90.1, which we observed in each example of pheromone increase.

**Supplementary Figure 4.**
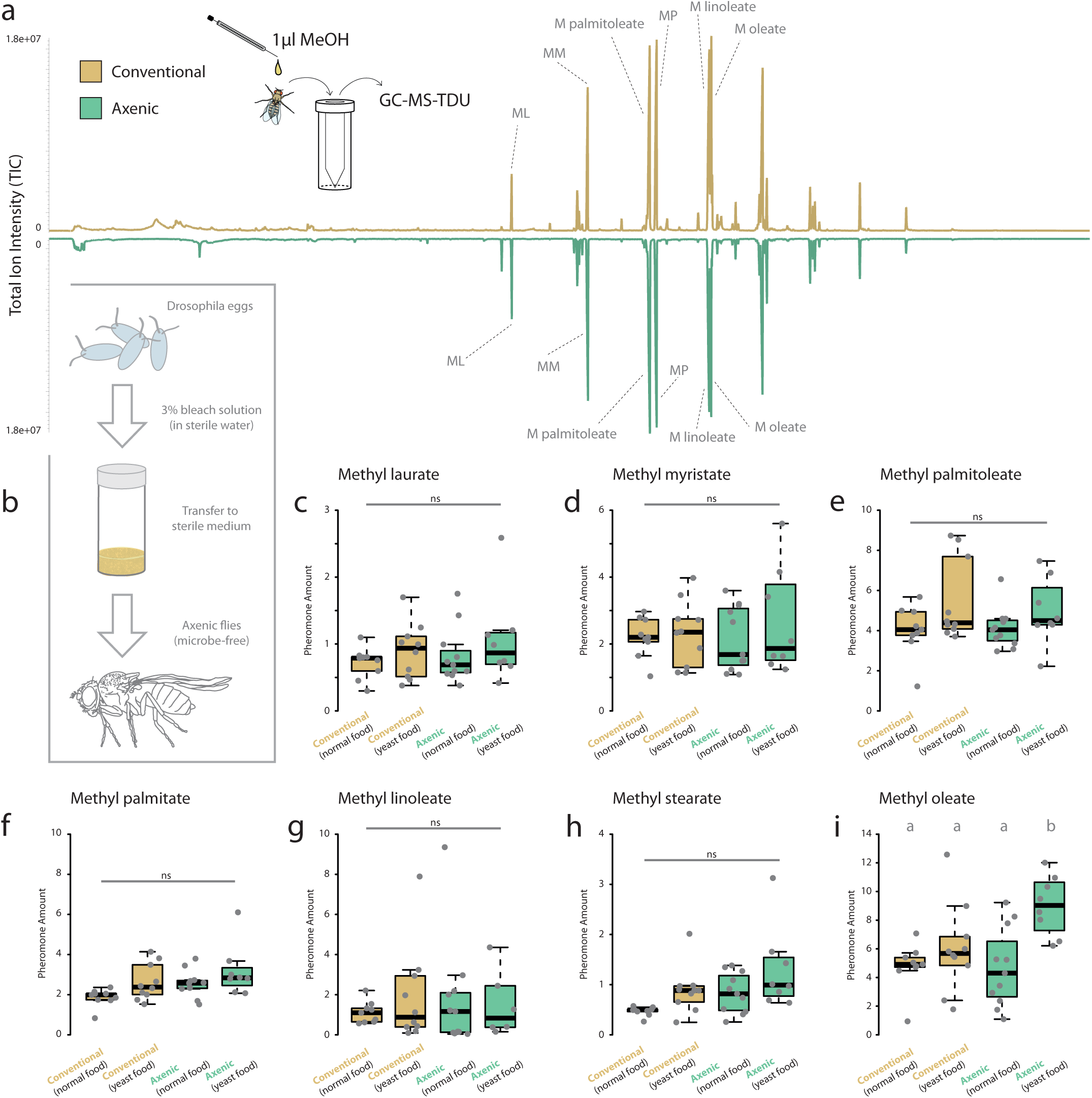
Pheromone comparisons between conventional and axenic Drosophila strains. (a) TDU-GC-MS analyses of odor profile from single adult flies exposed to droplet of methanol. Here we did not observe any significant differences in odor profiles between those flies grown with microbial symbionts (orange) and those without (green). (b) Diagram of methodology for axenic fly generation. Additional information available within the methods section. (c-i) Comparisons of average pheromone levels for Drosophila grown with natural microbial symbionts (orange) and those without (green; axenic). We only note a single difference, within methyl oleate, and only for axenic flies grown on food with (non-living) yeast extract within their media (i).

**Supplementary Figure 5.**
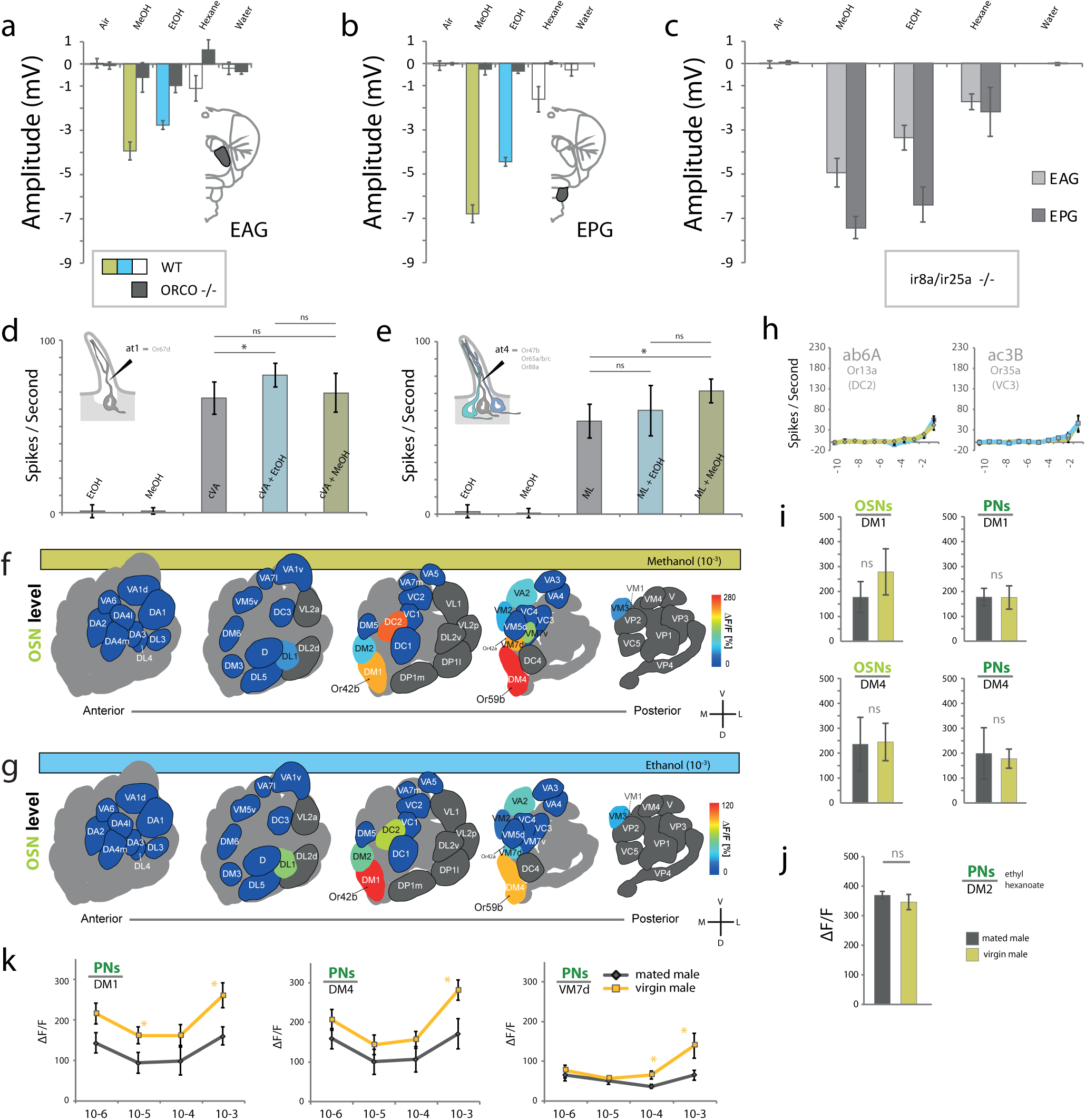
EAG, EPG, SSR and co-receptor mutants. (a) Electroantennogram, showing that the antenna can strongly detect both alcohols, but not for ORCO mutants. (b) Electropalpogram, showing that the palps of wildtype flies can detect both ethanol and methanol, but not for ORCO mutants. (c) EAG and EPG recordings towards odorants using ir8a/ir25a double mutant for IR co-receptors. (d) SSR recordings from at1 sensillum with alcohol exposure. (e) SSR recordings from at4 sensillum with alcohol exposure. (f,g) Antennal lobe (AL) diagram showing odor-induced calcium responses towards methanol and ethanol (10-3) within the olfactory sensory neurons (OSNs). (h) Dose response for SSR. (i) Odor-induced fluorescent activity towards ethanol within PNs. (j) PN response to a non-alcohol odorant across physiological states, showing the specificity of changes for alcohol circuits. (k) Dose response curves for OSN and PN towards methanol across mated and virgin males.

**Supplementary Figure 6.**
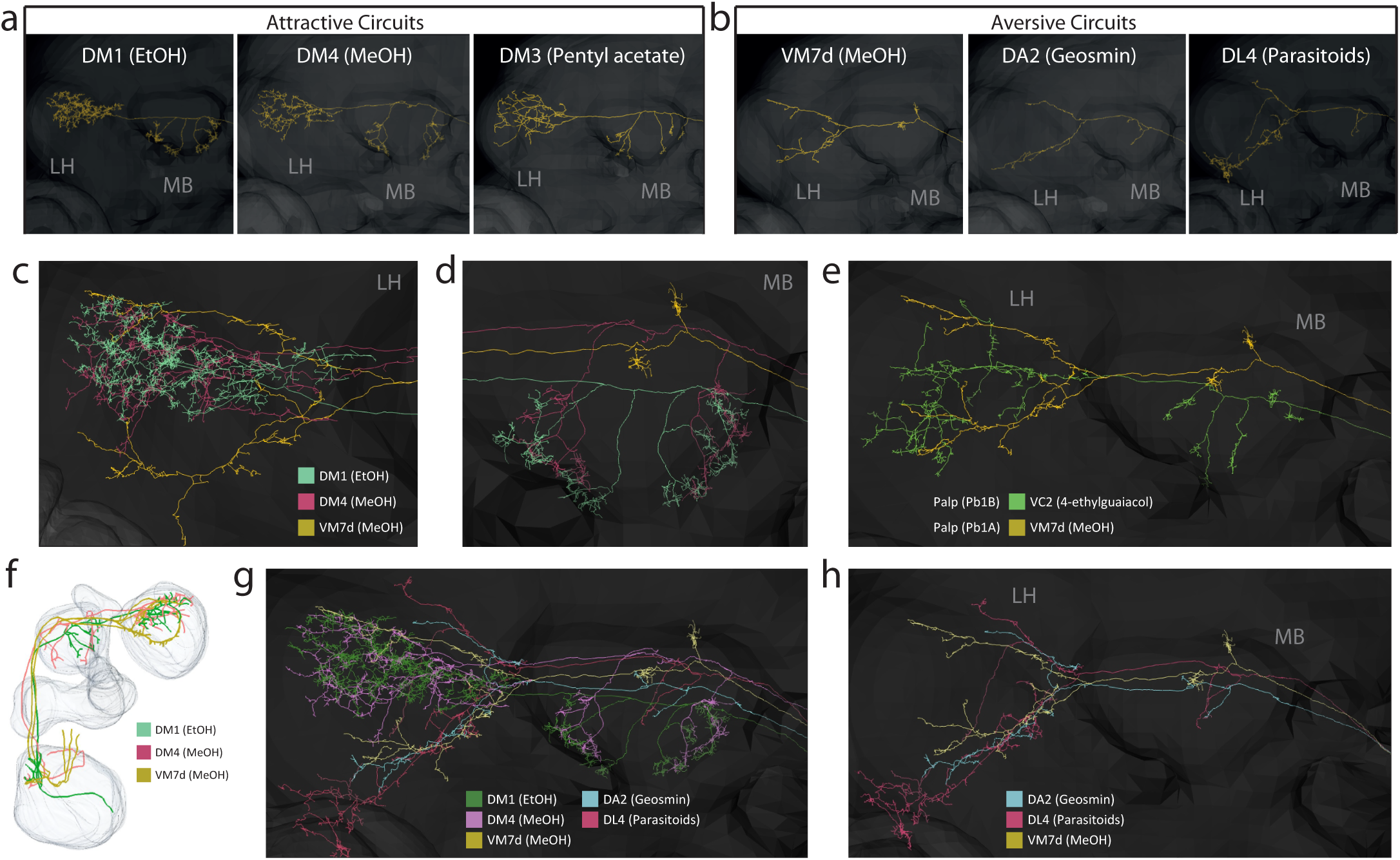
Neural reconstruction and circuitry related to alcohols. Single projection neuron (PN) reconstructions from mushroom body (MB) into the lateral horn (LH) 24–26. Upper zone of LH is associated with innate attractive behavior, while the lower portion is attributed to innate aversion. (a) Three single circuits associated with attractive behaviors. (b) Three single circuits associated with aversion. (c) Neural tracing of all three alcohol-related circuits within the LH, highlighting the difference between VM7d (yellow) and the two attractive circuits, DM1 and DM4. (d) Neural tracing of all three alcohol-related circuits within the MB, highlighting the difference between VM7d and the two attractive circuits, DM1 and DM4. (e) PN traces for both neurons extending from the pb1 sensillum showing that VM7d does not differ due to being from the palp rather than antenna. (f) PN reconstructions from AL through the mushroom body (MB) and into the lateral horn (LH). Note that VM7d (aversion) has three PNs while the attractive circuits (DM1, DM4) have only one. (g) All examined alcohol-related circuits, overlaid with known aversive tracts (DA2, DL4), note also the high number of terminals in MB for attraction but not aversive circuits. (h) VM7d circuit overlaid with known aversive tracts, showing high degree of overlap in LH as well as reduced MB branches.

**Supplementary Figure 7.**
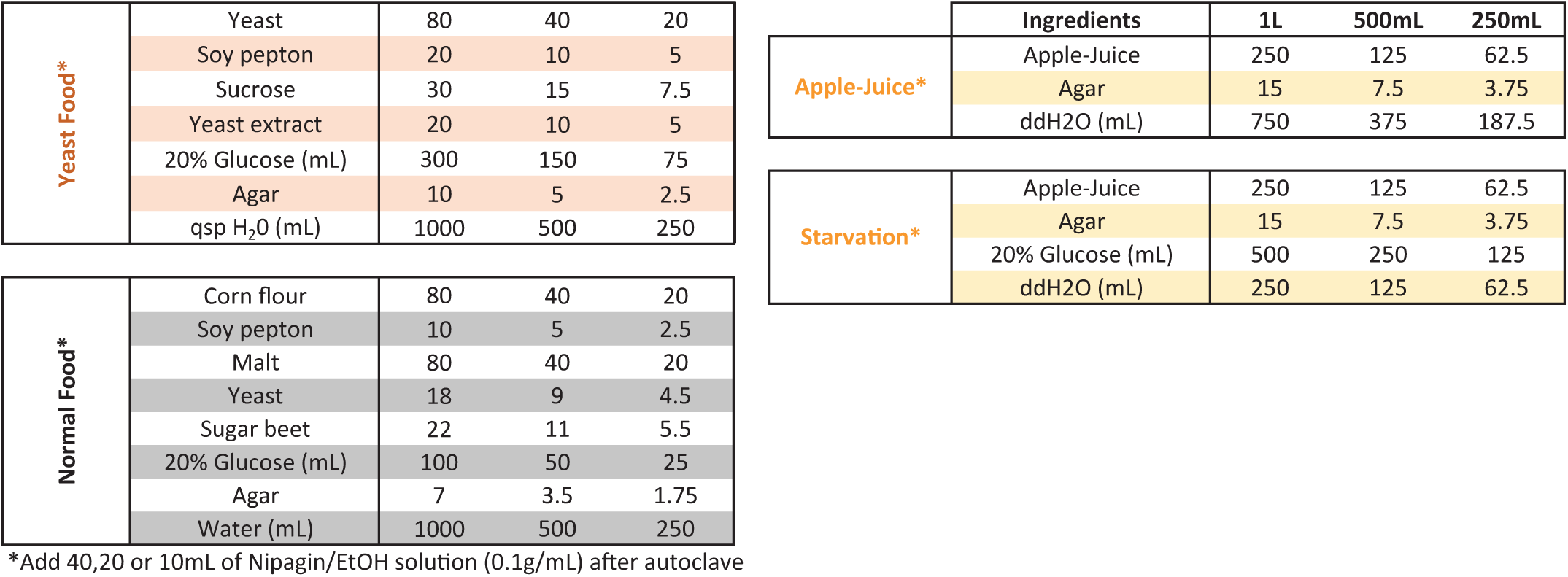
Diet and ingredients for axenic and gnotobiotic fly comparisons. Details are given for food types and media used for the generation and rearing of axenic and gnotobiotc flies 32,33. Additional information available within the methods section.

